# Three distinct neural mechanisms support movement-induced analgesia

**DOI:** 10.1101/2020.05.14.097261

**Authors:** Xuejing Lu, Xinru Yao, William Forde Thompson, Li Hu

## Abstract

Pain is essential for our survival by protecting us from severe injuries. Pain signals may be exacerbated by continued physical activities but can also be interrupted or over-ridden by physical movements, a process called movement-induced analgesia. A number of neural mechanisms have been proposed to account for this effect, including the reafference principle, the gate control theory of pain, and the top-down psychological modulation. Given that the analgesic effects of these mechanisms are temporally overlapping, it is unclear whether movement-induced analgesia results from a single neural mechanism or the joint action of multiple neural mechanisms. To address this question, we conducted five experiments on 130 healthy human subjects. First, the frequency of hand shaking was manipulated in order to quantify the relationship between the strength of the voluntary movement and the analgesic effect. Second, the temporal delay (between hand shaking and nociceptive laser stimuli) and the stimulated side (nociceptive laser stimuli were delivered on the hand ipsilateral or contralateral to the shaken one) were manipulated to quantify the temporal and spatial characteristics of the analgesic effect induced by voluntary movement. Combining psychophysics and electroencephalographic recordings, we demonstrated that movement-induced analgesia is a result of the joint action of multiple neural mechanisms. This investigation is the first to disentangle the distinct contributions of different neural mechanisms to the analgesic effect of voluntary movement. These findings extend our understanding of sensory attenuation arising from voluntary movement and may prove instrumental in the development of new strategies in pain management.

## Introduction

Pain sensations have a basic biological function: to direct an organism’s attention to injury and to protect it from further injury by triggering or inhibiting movement. The relationship between pain and movement is complex. When we touch a hot saucepan, we recoil in pain. When we twist an ankle, we avoid movement of that body part. Such responses have protective functions and instantiate adaptations to evolutionary selection pressures (1-3). However, there are circumstances in which pain signals appear to be inhibited in spite of continued use of an injured body part.

Movement-based analgesia refers to the process by which pain signals are interrupted or over-ridden by physical movement. There are countless examples of the apparent inhibition of pain signals in sports. Bert Trautmann, a German professional footballer, continued playing in the 1956 FA Cup Final despite having a severe injury to his neck, which was later determined to be broken. Similarly, Shun Fujimoto, a Japanese gymnast, smashed his knee during a dismount and yet continued to complete the pommel horse and the rings event. Such cases illustrate the powerful mechanisms that function to interrupt the transmission of pain signals. In addition to distraction, these mechanisms may be deployed through feedback from the physical movement itself, a phenomenon associated with sensory gating or sensory attenuation (4-6).

The sensory attenuation of afferent inputs (e.g., nociceptive inputs) induced by voluntary movement is conventionally held to prevent cognitive overload during movement execution and to free limited cognitive resources to attend to unpredicted, external events in the environment (7). This function is crucial as it enables us to dissociate afferents related to our own movements from those arising from the external environment (8), thus maintaining perceptual stability and guiding our behaviors (9). The underlying neural mechanism of sensory attenuation during voluntary movement fits well with a conceptual model, suggesting that a copy of the expected sensory results of a motor command or “an efference copy” is generated to anticipate sensory feedback from the movement (i.e., reafference) (10). When the reafference and the efference copy of the original motor command are of equal magnitude, reafferent information is eliminated from the sensory signal, and no sensory information is transmitted to the next levels of processing (11, 12). In addition to this reafference principle, the sensory attenuation of afferent inputs (especially nociceptive inputs) during voluntary movement can also be explained by other neural mechanisms. First, the gate control theory of pain holds that voluntary movement and its somatosensory feedback should activate large-diameter Aα and Aβ fibers, which inhibit the nociceptive volley transmitted via small-diameter Aδ and C fibers innervating spatially adjacent skin areas (13, 14). Second, sensory attenuation during voluntary movement may also occur through a top-down supraspinal descending inhibition mechanism, for example, arising from the anticipation of afferent inputs (15, 16).

The analgesic effect of voluntary movement during both movement preparation and execution is well documented (17-19). For instance, subjective pain perception ratings and brain responses elicited by nociceptive laser stimuli were significantly attenuated by the preparation of voluntary movement (20) and by moving the limb where laser stimuli were applied (21, 22). However, because the analgesic effects associated with different neural mechanisms are temporally overlapping, existing studies are unable to determine whether movement-induced analgesia results from a single neural mechanism or the joint action of multiple neural mechanisms. Isolating the independent and combined benefits of different analgesic mechanisms remains one of the most significant challenges in this emerging field.

One strategy for disentangling the effects of multiple mechanisms is to examine the differential time courses and spatial extensions of coexistent neural mechanisms. First, according to the reafference principle, the analgesic effect should be confined to spatially adjacent skin areas innervated by the reafferent nerves and should diminish with the termination of the motor command, thus producing a spatially localized, transient and reversible analgesic effect (5, 10). Second, in line with the gate control theory of pain, the analgesic effect should also be restricted to the innervating spatially adjacent skin areas, but persist after the termination of the somatosensory feedback, thus producing a spatially localized but sustained analgesic effect (14, 23). Third, the supraspinal descending inhibition mechanism arising from anticipation or attention about future events or outcomes should produce a spatially diffuse and transient analgesic effect (24). To demonstrate the differential temporal and spatial analgesic effects predicted by the three possible pain inhibition mechanisms, a simplified scheme is illustrated in Fig. 1.

**Figure 1.**
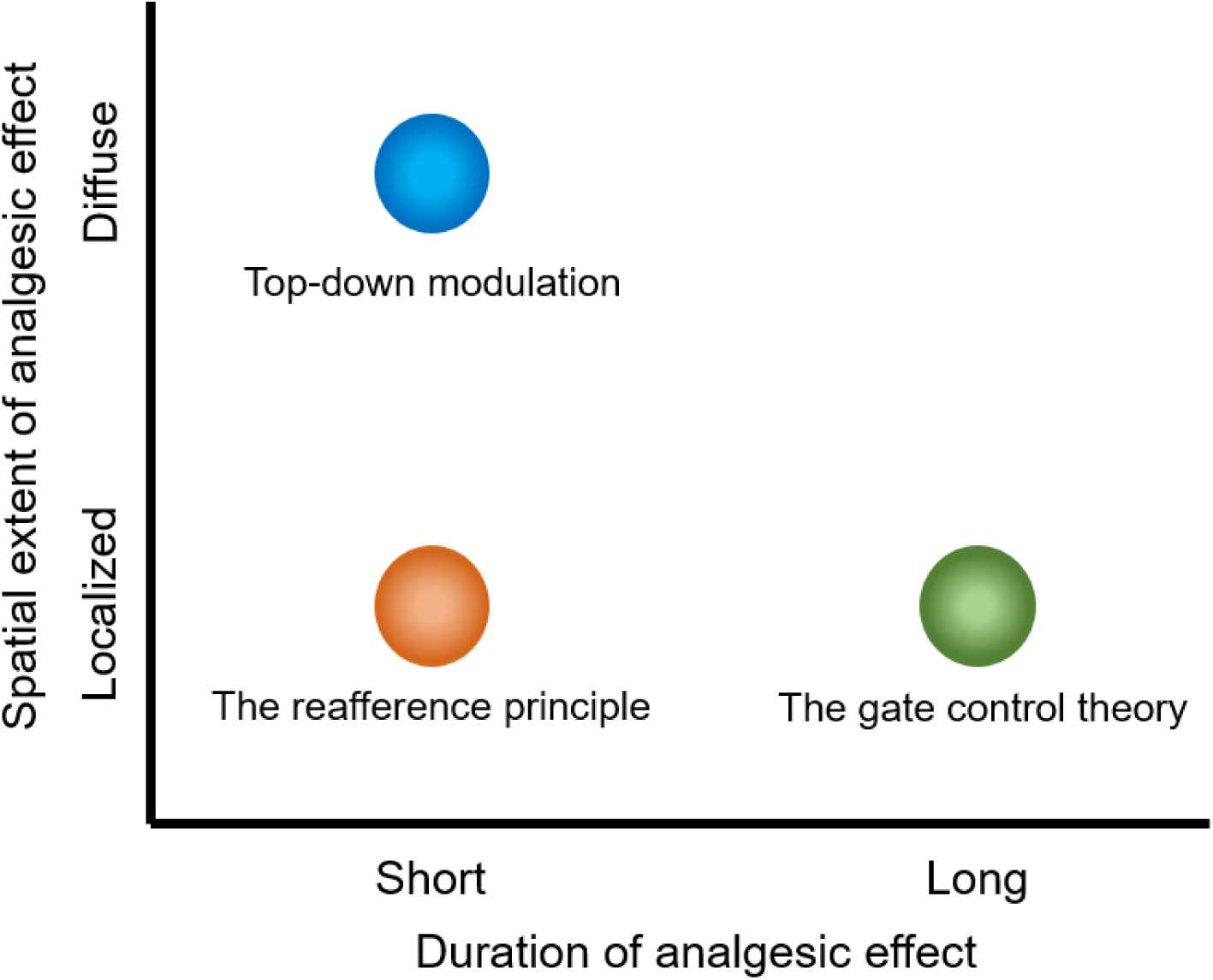
A simplified scheme of differential temporal and spatial analgesic effects predicted by three pain inhibition neural mechanisms. According to the reafference principle (the orange dot), the analgesic effect should be confined to spatially adjacent skin areas innervated by the reafferent nerves and should diminish with the termination of the motor command. The gate control theory of pain (the green dot) predicts an analgesic effect that should be restricted to the innervating spatially adjacent skin areas but persist after the termination of somatosensory feedback. The top-down descending inhibition mechanism (the blue dot) arising from anticipation or attention about future events or outcomes should produce a spatially diffuse and transient analgesic effect.

To isolate the respective analgesic effects of voluntary movement associated these different neural mechanisms, we conducted a series of experiments, in which subjects were instructed to shake their left or right hands for a short period of time (i.e., 5 s) before receiving nociceptive laser stimulus delivered on their left hands. Such manipulation allowed us to disentangle different neural mechanisms by applying nociceptive laser stimuli at different times and on different sites after the execution of voluntary movement. Before investigating the temporal and spatial characteristics of movement-induced analgesia, we established the relationship between voluntary movement and its analgesic effect by manipulating the strength of the movement.

## Results

We conducted five experiments on 130 healthy human subjects in total. High-power laser stimulation was applied to selectively excite cutaneous nociceptors and thereby elicit pure painful percepts without tactile sensations (25, 26). For all experiments, the energy of laser stimuli was individually determined by increasing stimulus energy in steps of 0.25 J, until a rating of 7 was obtained on a numerical rating scale ranging from 0 (no pain) to 10 (the worst pain imaginable). The analgesic effect was quantified by movement-induced changes of self-reports of pain intensity and unpleasantness, as well as electroencephalographic (EEG) responses. More details are described in **Materials and Methods**.

### The strength of voluntary movement modulates the analgesic effect

In Experiment 1, we collected psychophysical data from 15 healthy subjects (7 females, aged 22.1 ±1.9 years) to assess the relationship between the strength of voluntary movement and the analgesic effect by manipulating the frequency of hand shaking (Fig. 2*A*). Subjects were asked to rest their left hands on a table throughout the whole block (control condition) or shake their left hands for 5 s at approximately either 1 Hz (S1_low condition) or 5 Hz (S1_high condition). For each trial, nociceptive laser stimulus was delivered 1 s after the termination of hand movement, and subjects were instructed to provide self-reports of pain intensity and unpleasantness.

**Figure 2.**
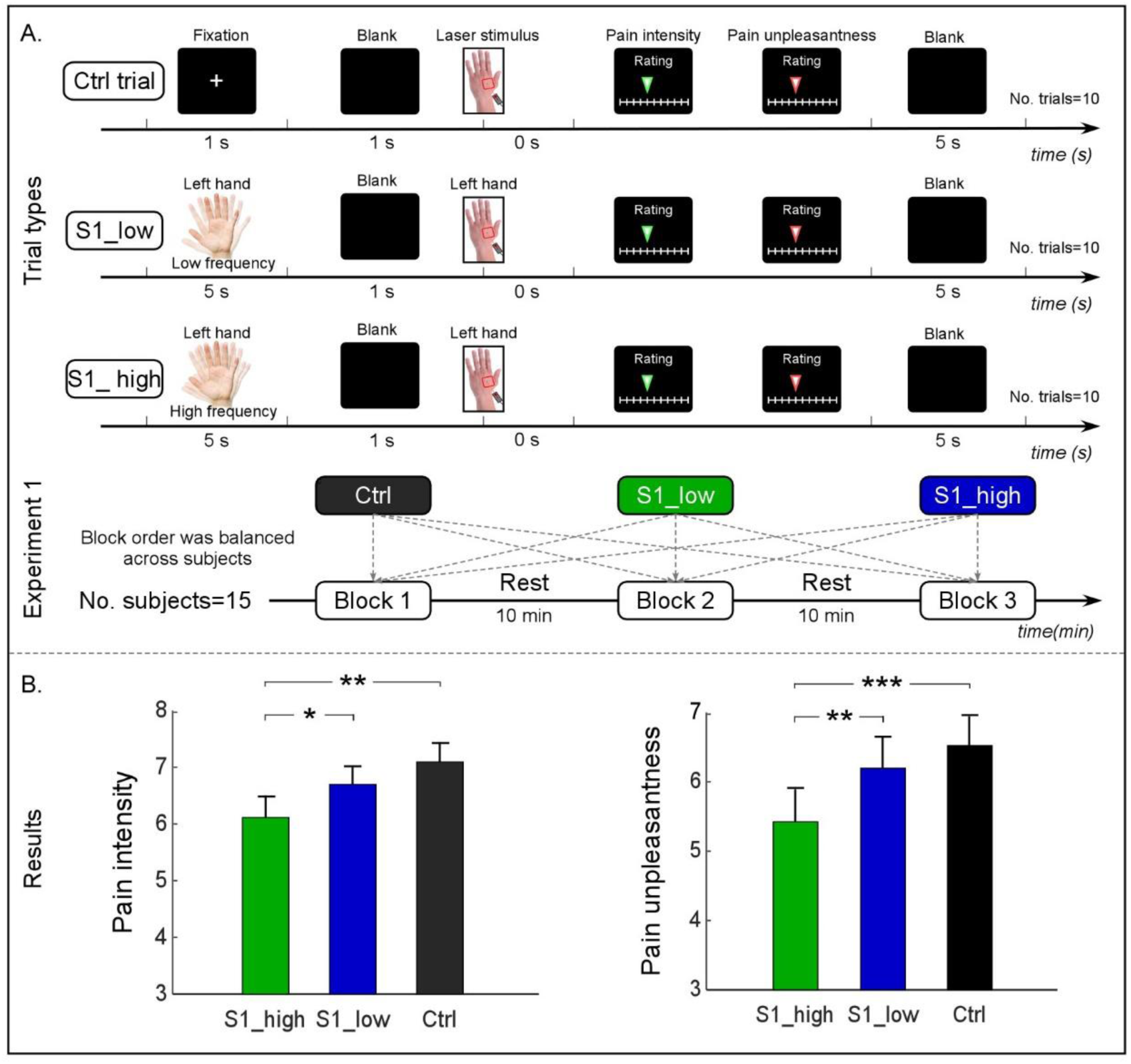
Experimental design and behavioral results of Experiment 1. (*A*) Experimental design. The experiment is composed of three blocks, each of which contains a distinct trial type (i.e., control, S1_low, or S1_high). Whereas subjects were asked to rest their left hands on a table in the control trial, they were asked to shake their left hands with a maximal range for 5 s at approximately 1 and 5 Hz in S1_low and S1_high trials, respectively. For each trial, 1 s after the termination of hand movement, nociceptive laser stimulus was delivered on the dorsum of their left hands. After each laser stimulus, subjects were instructed to rate the intensity and unpleasantness of the perceived pain using a 0–10 numerical rating scale. There were 10 trials in each block with a 5-s blank between two consecutive trials. After each block, there was a 10-min break. The block order was counterbalanced across subjects. (*B*) The comparison of subjective ratings of pain intensity (left) and unpleasantness (right) between different experimental conditions. *: *P* < 0.05; **: *P* < 0.01; ***: *P* < 0.001. Data are mean ± SEM.

One-way repeated-measures analysis of variance (ANOVA) showed strong evidence for a main effect of condition for both pain intensity (*F*_(2,28)_ = 9.70, *P* = 0.001, *η*_*p*_^*2*^ = 0.41) and unpleasantness (*F*_(2,28)_ = 11.89, *P* < 0.001, *η*_*p*_^*2*^ = 0.46) (Fig. 2*B* and *SI Appendix*, Table S1). Post hoc paired-sample t-tests showed that pain intensity and unpleasantness in the S1_high condition were significantly lower than those in the S1_low (intensity: *P* = 0.033; unpleasantness: *P* = 0.005) and control (intensity: *P* = 0.004; unpleasantness: *P* = 0.001) conditions. These results suggest that the more vigorous the movements are (i.e., a higher frequency of hand shaking), the stronger the analgesic effect of voluntary movement is.

### The temporal characteristics of the analgesic effect of voluntary movement

In Experiments 2 and 3, we collected self-reports of pain perception following nociceptive laser stimuli from 49 healthy subjects (Experiment 2: 24 subjects, 14 females, aged 21.9 ± 1.7 years; Experiment 3: 25 subjects, 14 females, aged 22.5 ± 2.3 years). Both experiments aimed to explore the temporal characteristics of the analgesic effect of voluntary movement by manipulating the temporal delay between hand shaking and nociceptive laser stimuli (short delays in Experiment 2 and long delays in Experiment 3).

In Experiment 2, subjects were asked to shake their left hands for 5 s at approximately 5 Hz, and nociceptive laser stimulus was delivered 1 s or 5 s after the termination of hand movement for the S1 and S5 conditions, respectively (Fig. 3*A* & 3*B*). One-way repeated-measures ANOVA showed strong evidence for a main effect of condition for both pain intensity (*F*_(2,46)_ = 16.80, *P* < 0.001, *η*_*p*_^*2*^ = 0.42) and unpleasantness (*F*_(2,46)_ = 18.95, *P* < 0.001, *η*_*p*_^*2*^ = 0.45) (Fig. 4*A*, and *SI Appendix*, Table S2). Post hoc paired-sample t-tests showed that pain intensity and unpleasantness in the control condition were significantly higher than those in the S1 and S5 conditions (all *P* < 0.001). No significant difference between the S1 and S5 conditions was observed for either pain intensity or unpleasantness (both *P* > 0.3). These results indicate that the analgesic effect of voluntary movement is strong when the temporal delay between hand shaking and nociceptive laser stimuli is short (e.g., less than 5 s).

**Figure 3.**
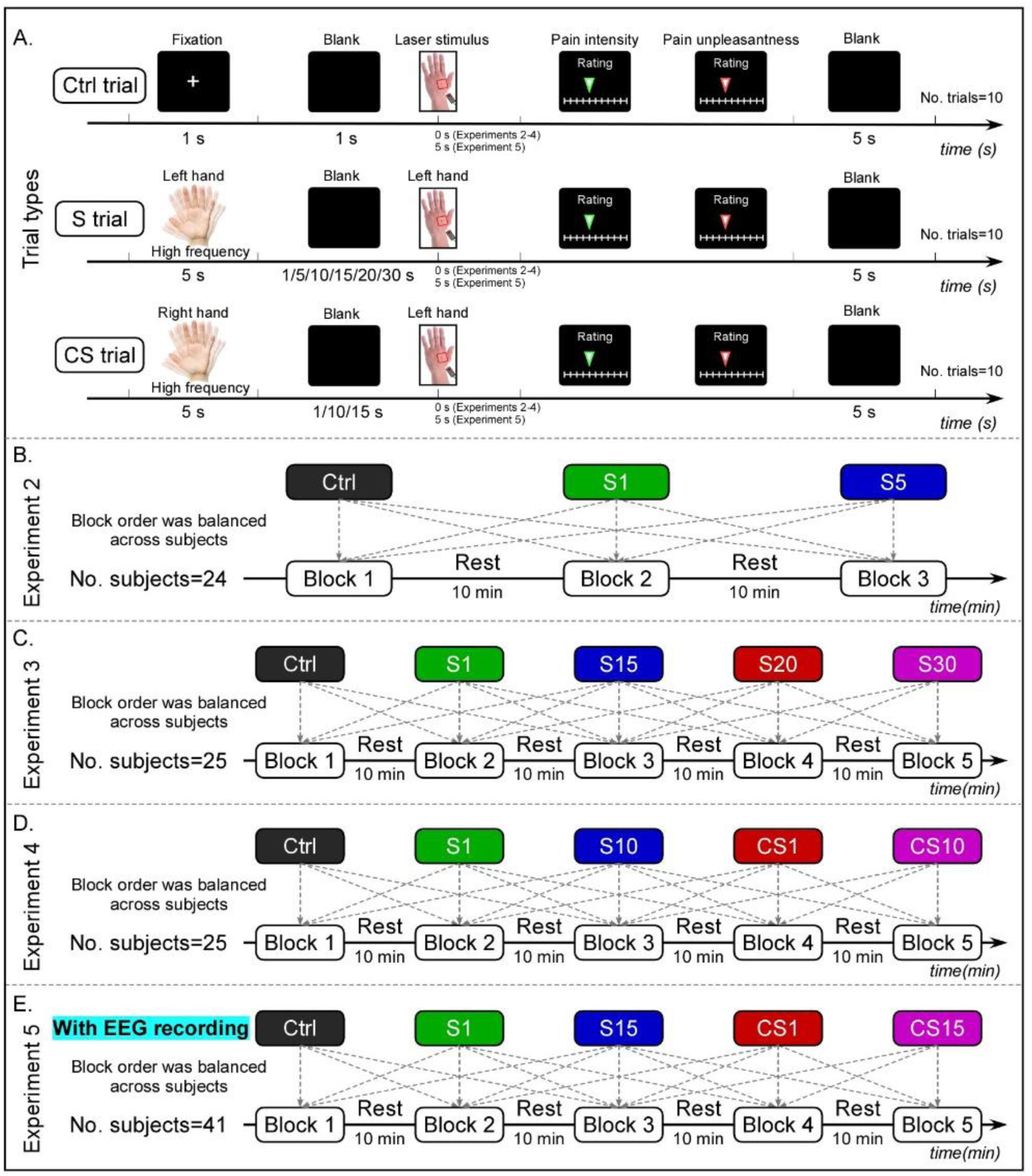
Experimental designs of Experiments 2-5. (*A*) Trial types. The control trials were identical to those in Experiment 1. The S trials were composed of S1, S5, S10, S15, S20, and S30 trials, for which subjects were asked to shake their left hands with a maximal range for 5 s at approximately 5 Hz, and nociceptive laser stimulus was delivered on the dorsum of their left hands 1 s, 5 s, 10 s, 15 s, 20 s, and 30 s after the termination of hand movement, respectively. The CS trials were composed of CS1, CS10, and CS15 trials, for which subjects were asked to shake their right hands with a maximal range for 5 s at approximately 5 Hz, and nociceptive laser stimulus was delivered on the dorsum of their left hands 1 s, 10 s, and 15 s after the termination of hand movement, respectively. For all trial types, subjects were instructed to rate the intensity and unpleasantness of the perceived pain using a 0–10 numerical rating scale after each laser stimulus. There was a 5-s blank between two consecutive trials. In addition, there was a 5-s interval between laser stimulus and the rating interface in Experiment 5 to ensure that the laser-evoked brain responses were not affected by rating procedures. (*B*) Experiment 2 is composed of three blocks, each of which contains a distinct trial type (i.e., control, S1, or S5). (*C*) Experiment 3 is composed of five blocks, each of which contains a distinct trial type (i.e., control, S1, S15, S20, or S30). (*D*) Experiment 4 is composed of five blocks, each of which contains a distinct trial type (i.e., control, S1, S10, CS1, or CS10). (*E*) Experiment 5 is composed of five blocks, each of which contains a distinct trial type (i.e., control, S1, S15, CS1, or CS15). EEG data were continuously recorded using 64 scalp electrodes in Experiment 5. For all experiments, there were 10 trials for each block, and the block order was counterbalanced across subjects. After each block, there was a 10-min break.

**Figure 4.**
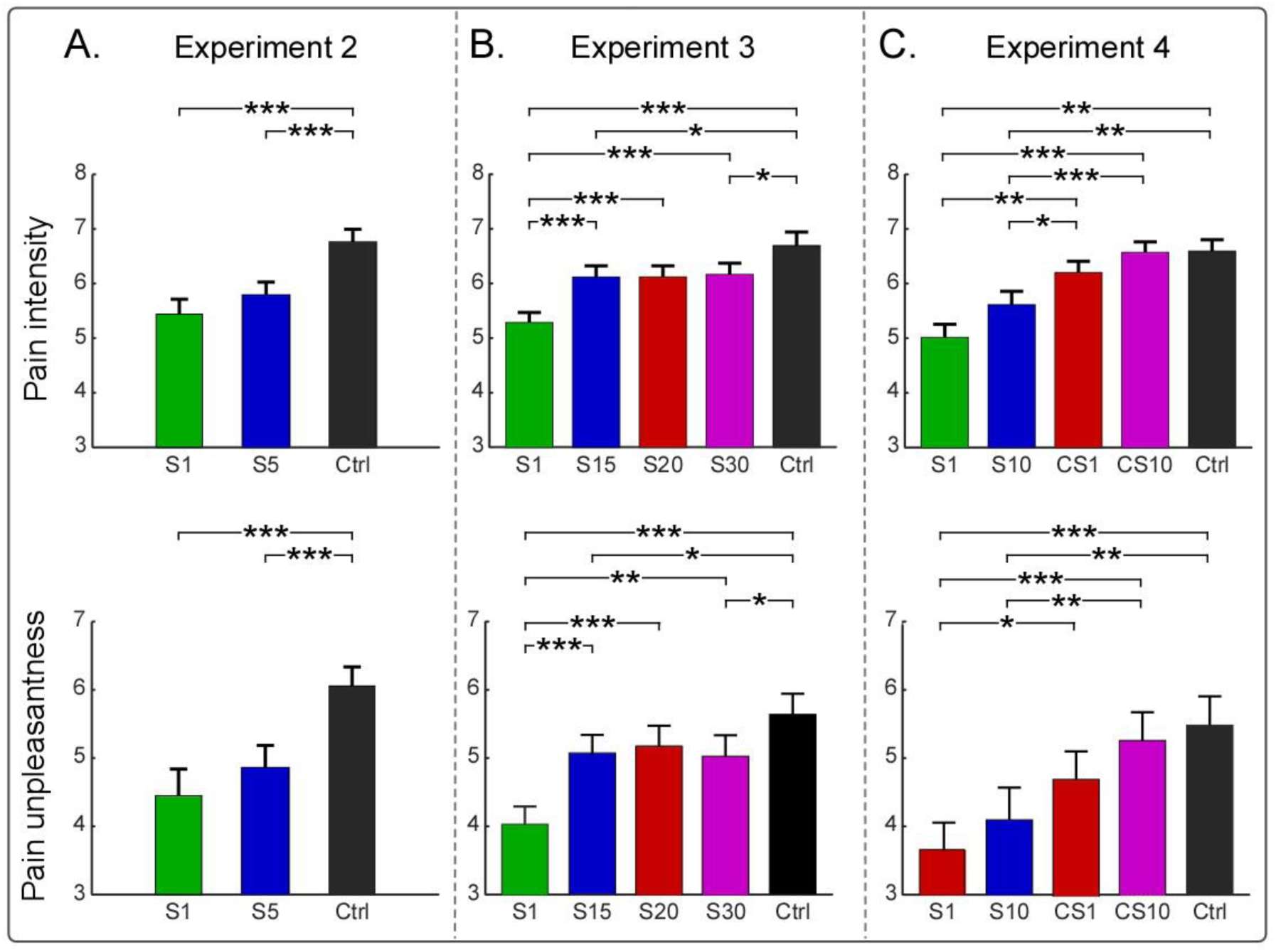
Behavioral results of Experiments 2-4. (*A-C*) The comparison of subjective ratings of pain intensity (top) and unpleasantness (bottom) between different experimental conditions in Experiments 2-4. *: *P* < 0.05; **: *P* < 0.01; ***: *P* < 0.001. Data are mean ± SEM.

In Experiment 3, the temporal delay between hand shaking and nociceptive laser stimuli was 1 s, 15 s, 20 s, and 30 s for S1, S15, S20, and S30 conditions, respectively (Fig. 3*A* & 3*C*). One-way repeated-measures ANOVA showed strong evidence for a main effect of condition for both pain intensity (*F*_(4,96)_ = 19.88, *P* < 0.001, *η*_*p*_^*2*^ = 0.45) and unpleasantness (*F*_(4,96)_ = 16.78, *P* < 0.001, *η*_*p*_^*2*^ = 0.41) (Fig. 4*B* and *SI Appendix*, Table S3). Post hoc paired-sample t-tests showed that pain intensity and unpleasantness in the control condition were significantly higher than those in the S1 (both *P* < 0.001), S15 (intensity: *P* = 0.019; unpleasantness: *P* = 0.048), and S30 (intensity: *P* = 0.036; unpleasantness: *P* = 0.011) conditions. Pain intensity in the control condition was marginally higher than that in the S20 condition (*P* = 0.051). In addition, pain intensity and unpleasantness in the S1 condition were significantly lower than those in the S15 (both *P* < 0.001), S20 (intensity: *P* < 0.001; unpleasantness: *P* = 0.003), and S30 (both *P* < 0.001) conditions. No significant difference was observed for either pain intensity or unpleasantness in the S15, S20, and S30 conditions (all *P* > 0.9). These results indicate that when the temporal delay between hand shaking and nociceptive laser stimuli is long (i.e., from 15 s to 30 s), the analgesic effect of voluntary movement is evident but obviously weaker than the short temporal delay (i.e., 1 and 5 s).

### The spatial characteristics of the analgesic effect of voluntary movement

In Experiments 4 and 5, we collected self-reports of pain perception following nociceptive laser stimuli from 66 healthy subjects (Experiment 4: 25 subjects, 14 females, aged 23.6 ± 2.4 years; Experiment 5: 41 subjects, 24 females, aged 22.7 ± 1.9 years). Both experiments aimed to investigate the spatial characteristics of the analgesic effect of voluntary movement by manipulating the stimulated side (i.e., nociceptive laser stimuli were delivered on the hand ipsilateral or contralateral to the shaken one). Whereas only behavioral data were collected in Experiment 4, we coupled psychophysics with high-density EEG recordings in Experiment 5.

In Experiment 4, nociceptive laser stimulus was delivered on left hand dorsum 1 s or 10 s after the termination of left-hand movement for the S1 and S10 conditions, respectively. In contrast, nociceptive laser stimulus was delivered on left hand dorsum 1 s or 10 s after the termination of right-hand movement for the CS1 and CS10 conditions, respectively (Fig. 3*A* & 3*D*). One-way repeated-measures ANOVA showed strong evidence for a main effect of condition for both pain intensity (*F*_(4,96)_ = 18.32, *P* < 0.001, *η*_*p*_^*2*^ = 0.43) and unpleasantness (*F*_(4,96)_ = 13.00, *P* < 0.001, *η*_*p*_^*2*^ = 0.35) (Fig. 4*C* and *SI Appendix*, Table S4). Post hoc paired-sample t-tests showed that pain intensity and unpleasantness in the control condition were significantly higher than those in the S1 (both *P* ≤ 0.001) and S10 (intensity: *P* = 0.007; unpleasantness: *P* = 0.004) conditions, but not than those in the CS1 and CS10 conditions (all *P* ≥ 0.09). Pain intensity and unpleasantness in the S1 condition were significantly lower than those in the CS1 (intensity: *P* = 0.002; unpleasantness: *P* = 0.041) and CS10 (both *P* < 0.001) conditions. Pain intensity in the S10 condition was significantly lower than that in the CS1 (*P* = 0.035) and CS10 (*P* < 0.001) conditions, and pain unpleasantness in the S10 condition was significantly lower than that in the CS10 condition (*P* = 0.008). These results indicate that the analgesic effect of voluntary movement is greatly influenced by the stimulated side: the analgesic effect is much stronger when nociceptive laser stimuli are delivered on the same hand with voluntary movement than on the other hand. Importantly, there was a clear trend that self-reports in the CS1 condition were smaller than in the control condition (unpleasantness: uncorrected *P* = 0.009, Bonferroni-corrected *P* = 0.09), suggesting a possible spatially diffuse and transient analgesic effect.

To assess whether the spatially diffuse and transient analgesic effect is evident, we collected psychophysical data from an additional group of 41 subjects in Experiment 5. The experiment design was similar with that of Experiment 4, except that instead of the S10 and CS10 conditions, the S15 and CS15 conditions were included in Experiment 5, in which nociceptive laser stimulus was delivered 15 s after the termination of voluntary movement (Fig. 3*A* & 3*E*). One-way repeated-measures ANOVA showed strong evidence for a main effect of condition for both pain intensity (*F*_(4,160)_ = 51.92, *P* < 0.001, *η*_*p*_^*2*^ = 0.57) and unpleasantness (*F*_(4,160)_ = 37.32, *P* < 0.001, *η*_*p*_^*2*^ = 0.48) (Fig. 5*A*). Post hoc paired-sample t-tests showed that pain intensity and unpleasantness in the control condition were significantly higher than those in the S1 (both *P* < 0.001), S15 (both *P* < 0.001), and CS1 (intensity: *P* = 0.029; unpleasantness: *P* = 0.018) conditions, but not than those in the CS15 condition (both *P* > 0.05). Pain intensity and unpleasantness in the S1 condition were significantly lower than those in the S15, CS1, and CS15 conditions (all *P* < 0.001). Pain intensity and unpleasantness in the S15 condition were significantly lower than those in the CS1 (intensity: *P* = 0.001; unpleasantness: *P* = 0.021) and CS15 (intensity: *P* = 0.002; unpleasantness: *P* = 0.007) conditions. Similar to Experiment 4, these results suggest that the analgesic effect of voluntary movement is greatly influenced by the stimulated side. Importantly, there is a clear analgesic effect in the CS1 condition, demonstrating the existence of the spatially diffuse and transient analgesic effect of voluntary movement. Moreover, different from the observation in Experiment 4 where self-reports of pain perception between the S1 and S10 conditions were not significant, we found that self-reports between the S1 and S15 conditions were significantly different in Experiment 5 (in line with the observation in Experiment 3). This observation suggests that the analgesic effect is stronger when the temporal delay was shorter than 10 s, but much weaker when the temporal delay is longer than 15 s.

**Figure 5.**
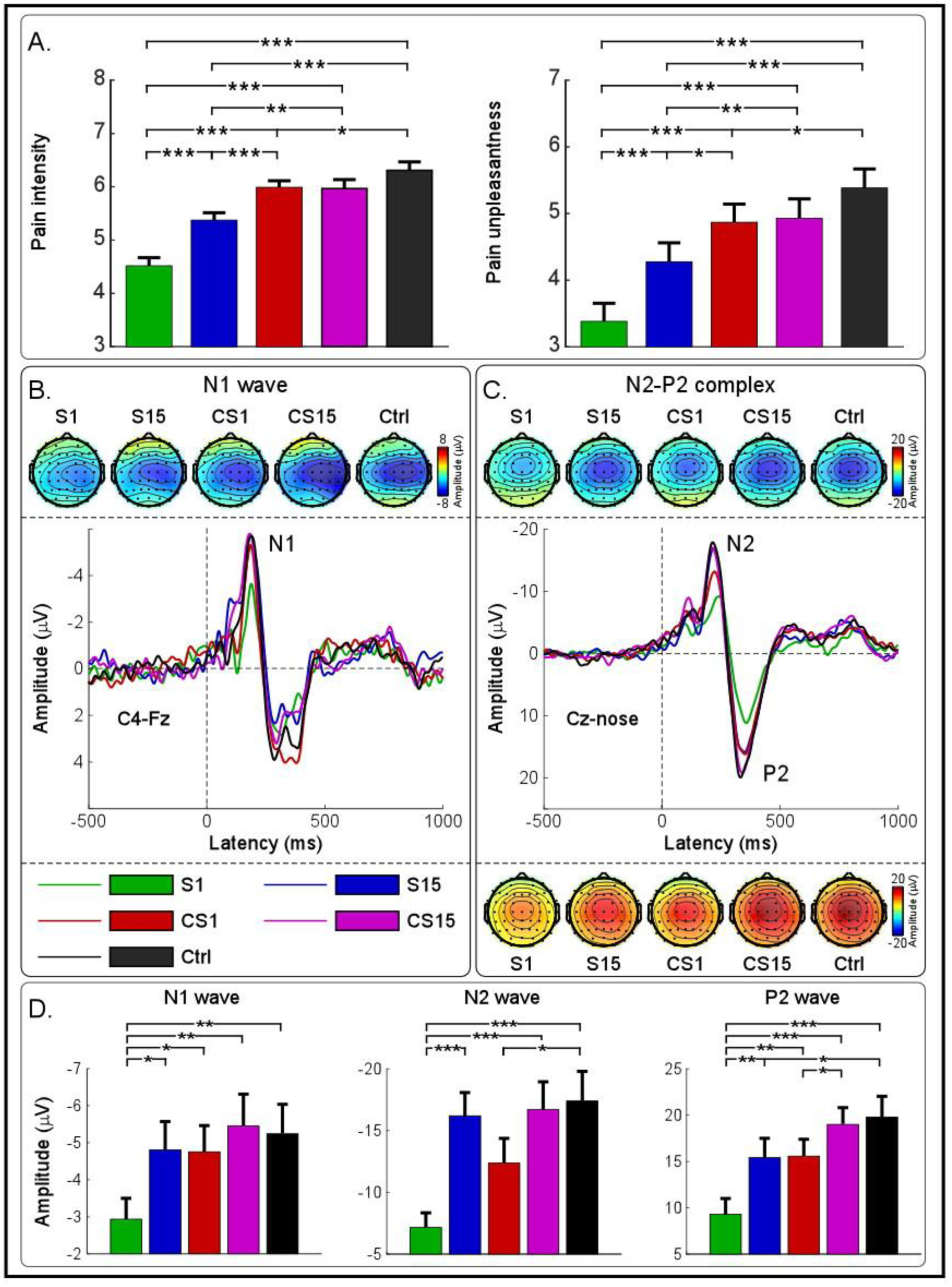
Behavioral and electrophysiological results of Experiment 5. (*A*) The comparison of subjective ratings of pain intensity (left) and unpleasantness (right) between different experimental conditions. (*B-C*) Group-level LEP waveforms and scalp topographies of the N1, N2, and P2 waves are displayed for each experimental condition. (*D*) The comparison of the N1, N2, and P2 amplitudes between different experimental conditions. *: *P* < 0.05; **: *P* < 0.01; ***: *P* < 0.001. Data are mean ± SEM.

### The modulation of voluntary movement on pain-related brain responses

Group-level laser-evoked brain potentials (LEPs) and scalp topographies of the N1, N2, and P2 waves are shown in Fig. 5*B* & 5*C*. In line with previous observations (25, 26), scalp topographies of the earliest activity in LEPs (i.e., the N1 wave) showed a clear negative maximum on the central electrodes overlying the hemisphere contralateral to the stimulated side. Scalp topographies of the N2 wave were maximal at the vertex and extended bilaterally toward temporal regions, and scalp topographies of the P2 wave were more centrally distributed.

There were clear modulations of voluntary movement on the N1, N2, and P2 waves, and one-way repeated-measures ANOVA showed strong evidence for a main effect of condition for the N1 (*F*_(4,160)_ = 5.98, *P* < 0.001, *η*_*p*_^*2*^ = 0.13), N2 (*F*_(4,160)_ = 10.17, *P* < 0.001, *η*_*p*_^*2*^ = 0.20), and P2 (*F*_(4,160)_ = 16.34, *P* < 0.001, *η*_*p*_^*2*^ = 0.29) amplitudes (Fig. 5*D* and *SI Appendix*, Table S5). Post hoc paired-sample t-tests showed that the N1 amplitude in the S1 condition was significantly lower than those in all other conditions (all *P* < 0.05), and no significant condition difference was observed between S15, CS1, CS15, and control conditions (all *P* > 0.05). Since the N1 wave represents somatosensory specific activities maximally reflecting the incoming nociceptive inputs (27), the N1 results suggest that in the S1 condition the nociceptive information has been greatly attenuated before reaching to the cerebral cortex.

The N2 amplitude in the S1 condition was significantly lower than those in the S15, CS15, and control conditions (all *P* < 0.001), but not than that in the CS1 condition (*P* = 0.084). Furthermore, the N2 amplitude in the CS1 condition was significantly lower than that in the control condition (*P* = 0.015). In other words, the N2 amplitude was reduced when the temporal delay between hand shaking and nociceptive laser stimuli was short (i.e., 1 s), regardless of whether laser stimuli were delivered on the same or the different hand with voluntary movement. Considering that the N2 wave is highly associated with attentional factors (28, 29), the modulation of voluntary movement on the N2 amplitude would reflect the top-down descending pain inhibition, thus providing additional evidence to support the existence of the spatially diffuse and transient analgesic effect of voluntary movement.

The P2 amplitude in the S1 condition was significantly lower than those in all other conditions (all *P* < 0.01). In addition, the P2 amplitude in the S15 condition was significantly lower than that in the control condition (*P* = 0.039), and marginally lower than that in the CS15 condition (uncorrected *P* = 0.0065, Bonferroni-corrected *P* = 0.065). The P2 amplitude in the CS1 condition was significantly lower than that in the CS15 condition (*P* = 0.025). These results suggest the coexistence of a spatially localized analgesic effect that may persist for a period of time (i.e., S15 condition), and a spatially diffuse and transient analgesic effect (i.e., CS1 condition).

### The modulation of voluntary movement on spontaneous EEG oscillations

Group-level spectra of spontaneous EEG oscillations, together with the statistical results, are shown in Fig. 6. Clear differences of spectral amplitude among experimental conditions were observed at alpha frequencies (8-12 Hz, Fig. 6*A*).

**Figure 6.**
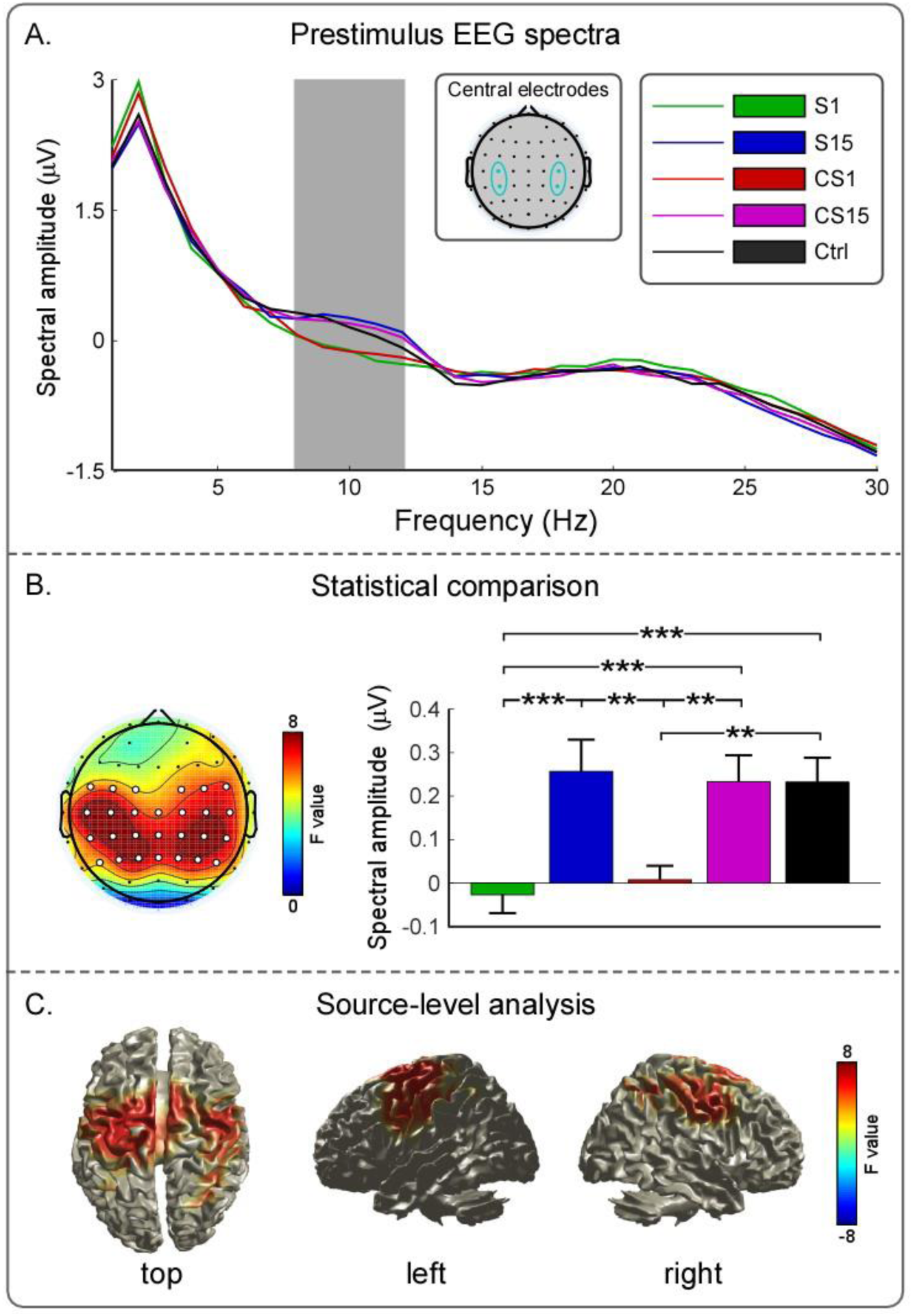
The effect of voluntary movement on spontaneous EEG oscillations (Experiment 5). (*A*) Group-level spectra of spontaneous EEG oscillations at bilateral central electrodes (C3, CP3, C4, and CP4) are displayed for each experimental condition. A clear difference of spectral amplitude was observed at alpha frequencies (8-12 Hz, gray area). (*B*) The comparison of spectral amplitude of alpha oscillations between different experimental conditions. **: *P* < 0.01; ***: *P* < 0.001. Data are mean ± SEM. The modulation of voluntary movement on alpha oscillations was maximal at bilateral central regions at the scalp level (electrodes with clear evidence for a main effect of condition are marked in white, *P* < 0.05, corrected using a false discovery rate [FDR] procedure). (*C*) The modulation of voluntary movement on alpha oscillations was maximal at bilateral primary sensorimotor cortices at the source level (brain regions with clear evidence for a main effect of condition are color-coded, FDR-corrected *P* < 0.05).

One-way repeated-measures ANOVA showed strong evidence for the main effect of condition for the spectral amplitude of alpha oscillations (*F*_(4,160)_ = 15.15, *P* < 0.001, *η*_*p*_^*2*^ = 0.28). Post-hoc paired-sample t-tests showed that the spectral amplitude of alpha oscillations in the S1 condition was significantly lower than those in the S15, CS15, and control conditions (all *P* < 0.001). In addition, the spectral amplitude of alpha oscillations in the CS1 condition was significantly lower than those in the S15 (*P* = 0.006), CS15 (*P* = 0.002), and control (*P* < 0.001) conditions (Fig. 6*B*). At the scalp level, the modulation of voluntary movement on alpha oscillations was maximal at bilateral central regions (Fig. 6*B*). At the source level, such modulation was observed at bilateral primary sensorimotor cortices (S1/M1, Fig. 6*C*). These results showed that voluntary movement induced a clear change in the spontaneous brain state, i.e., a decrease of the spectral amplitude of alpha oscillations in bilateral S1/M1, which could play an active role for the top-down inhibitory control of the nociceptive information (30, 31). Therefore, the modulation of voluntary movement on the functional state of bilateral S1/M1 provides an electrophysiological basis for the spatially diffuse and transient analgesic effect of voluntary movement.

## Discussion

In the present study, a series of experiments were conducted on 130 healthy subjects to disentangle the underlying mechanisms of movement-induced analgesia. First, the frequency of hand shaking was manipulated in Experiment 1, and we observed that more vigorous movements confer more significant analgesic benefits (Fig. 2*B*). Second, the temporal delay between hand shaking and nociceptive laser stimuli was manipulated in Experiments 2 and 3 to quantify the temporal characteristics of the analgesic effect. When the temporal delay was short (e.g., less than 15 s), the analgesic effect of voluntary movement was strong but decreased rapidly. When the temporal delay was long (e.g., from 15 s to 30 s), the analgesic effect was weak but sustained (Fig. 4*A* & 4*B*). These observations suggest the coexistence of two different analgesic effects: one is strong but transient, and the other is weak but sustained. Third, the stimulated side (nociceptive laser stimuli were delivered on the hand ipsilateral or contralateral to the shaken one) was manipulated in Experiments 4 and 5 to quantify the spatial characteristics of the analgesic effect. When nociceptive laser stimuli were delivered on the same hand with voluntary movement, the analgesic effect was much stronger than when they were delivered on the other hand. Additionally, if the temporal delay between hand shaking and nociceptive laser stimuli was very short (i.e., 1 s), there was a weak analgesic effect even when laser stimuli were delivered on the hand without voluntary movement (Figs. 4*C* & 5*A*). These observations suggest the existence of another analgesic effect, which is weak but spatially diffuse, besides the two spatially localized analgesic effects observed in Experiments 2 and 3.

In addition to the behavioral results, Experiment 5 provided electrophysiological evidence (i.e., the modulation of voluntary movement on the N1, N2, and P2 amplitudes, Fig. 5*D*) to support the coexistence of distinct analgesic effects of voluntary movement. Moreover, clear modulation of voluntary movement on spontaneous alpha oscillations was also observed in Experiment 5 (Fig. 6), suggesting changes of the functional state of bilateral primary sensorimotor cortices due to voluntary movement, which could play an active role for the top-down inhibitory control of the nociceptive information.

Altogether, coupled psychophysics with high-density EEG recordings, our study provides a comprehensive understanding of the temporal and spatial characteristics of the analgesic effect related to voluntary movement, revealing that movement-induced analgesia is a result of the joint action of multiple neural mechanisms. Our findings not only inform the field of pain neuroscience but may also help guide the development of non-pharmaceutical analgesic strategies in clinical practice.

### The analgesic effect associated with the reafference principle

The analgesic effect of voluntary movement was strongly modulated by the frequency of hand shaking. This effect rapidly decreased within a short time window (e.g., less than 15 s) and was largely confined to spatially adjacent skin areas of the innervated hand with voluntary movement (Fig. 4). These results revealed that voluntary movement confers a spatially localized, transient and reversible analgesic effect. These analgesic characteristics are predicted by the reafference principle; that is, when a motor command is generated, an efference copy of this command is utilized to predict the sensory consequence of the action (11, 12). If the reafference due to self-generated action and an efference copy of the initial motor command are of equal magnitude, they cancel out (32). The generation of such inhibition can be attributed to the descending motor commands, rather than the peripheral feedback from the motion itself (4, 18), given that inhibitions occurred preferentially during active movement, and were observed before the recording of the electromyographic activity (19).

By recording the electrophysiological activity from the radial nerve of awake, behaving monkeys, Seki, Perlmutter and Fetz (33) found that cutaneous afferent inputs recorded from interneurons at the spinal cord were inhibited presynaptically during voluntary movement, thus effectively reducing synaptic transmission at the initial synapse. This observation suggests that the analgesic effect associated with the reafference principle is spatially localized. In addition, our brain continuously predicts the sensory consequences of its own body movements in an active fashion (34, 35) to dynamically minimize prediction errors (36). This essential brain function, along with the dynamic property of the modulation of descending motor commands (7, 37), accounts for why the reduction of sensory transmission caused by voluntary movement is transient and reversible.

### The analgesic effect associated with the gate control theory of pain

The analgesic effect of voluntary movement ipsilateral to nociceptive laser stimuli was sustained within a long time window, reaching a plateau with no further decline when the temporal delay between hand shaking and nociceptive laser stimuli was relatively long (i.e., from 15 to 30 s, Fig. 4*B*). However, within this time window, no analgesic effect was observed if nociceptive laser stimuli were delivered on the hand without voluntary movement (i.e., the CS15 condition in Experiment 5, Fig. 5). These findings suggest a spatially localized but sustained analgesic effect, which cannot be explained by the reafference principle, as the motor commands should be diminished with the termination of voluntary movement. Instead, they suggest an alternative mechanism, predicted by the gate control theory of pain. Indeed, hand shaking and its somatosensory feedback should activate large-diameter Aα and Aβ fibers, which inhibit the nociceptive volley transmitted via small-diameter Aδ and C fibers innervating spatially adjacent skin areas (13, 23).

This explanation assumes the same neural mechanism of tactile-induced analgesia, in which the activation of tactile afferent nerve fibers (e.g., large-diameter Aβ fibers) inhibits the transmission of nociceptive inputs (38-40). Such an analgesic effect was maximal when tactile inputs were delivered homotopically to the same body territory of pain (23). This phenomenon fits well with our observation that the sustained analgesic effect was only observed at the stimulated side ipsilateral to hand movement. In addition, it has been demonstrated that tactile-induced analgesia can persist for up to hours in non-human animals (41, 42) and humans (23), which is consistent with our observation that the analgesic effect persisted over a period of time after the execution of voluntary movement (e.g., up to 30 s in our case, Fig. 4*B*).

### The analgesic effect associated with the top-down psychological modulation

The combination of both the reafference principle and the gate control theory of pain is sufficient to explain most findings of the present study, with the exception of the weak spatially diffuse analgesic effect; that is, an analgesic effect was observed when there was a very short temporal delay between hand shaking and nociceptive laser stimuli delivered on the hand without voluntary movement (i.e., the CS1 condition in Experiments 4 and 5). Importantly, this weak spatially diffuse analgesic effect was accompanied by the changes of spectral amplitude of spontaneous alpha oscillations at bilateral central regions (Fig. 6). Specifically, voluntary movement induced a decrease of the spectral amplitude of alpha oscillations in bilateral S1/M1, and such modulation provides an electrophysiological basis for the top-down psychological modulation of the nociceptive information (30, 31).

The functional state of the cerebral cortex can be partly indexed by spontaneous alpha oscillations, which would be dynamically modulated by hand movement and somatosensory feedback (43). For instance, alpha oscillations at bilateral central regions, originated from primary sensorimotor cortices, have been demonstrated to be desynchronized by voluntary movement and somatosensory stimulation, which is characterized as the decreased amplitude of the oscillations (44). Such a decrease was encoded by temporal anticipation, which subserved as a plausible mechanism for biasing subsequent perception and brain responses (45). This explanation is in line with our observation that, compared to the long temporal delays (e.g., 10 s and 15 s), subjects tended to have higher temporal anticipation for the forthcoming stimuli when the temporal delay between hand shaking and nociceptive laser stimuli was shorter (e.g., 1 s). Importantly, the modulation effect of anticipation on pain perception and brain responses is transient and reversible (46), which provided an apparent reason why the weak spatially diffuse analgesic effect was only observed following a short temporal delay. Indeed, our results cannot exclude another possible explanation for the weak spatially diffuse analgesic effect, which is attention modulation, for two reasons. First, drawing attention away from nociceptive laser stimuli during and after movement execution should reduce pain regardless of where the stimuli were delivered (28, 47). Second, the changes of alpha oscillations, which is highly associated with the weak spatially diffuse analgesic effect, could also be influenced by the attention modulation (48-50).

In summary, our findings suggest that movement can interrupt or override the pain signals if there are urgent demands on an organism. Understanding and isolating the independent and combined benefits of different analgesic mechanisms remains one of the most fundamental challenges in pain research. The present study was the first investigation to disentangle the distinct contributions of multiple neural mechanisms underlying the movement-induced analgesia. Specifically, movement-induced analgesia is a result of the joint action of multiple neural mechanisms, including but not limited to the reafference principle, the gate control theory of pain, and the top-down psychological modulation. The findings also extend our understandings of sensory attenuation arising from voluntary movement by generalizing the phenomenon from movement preparation and execution stages to the stage after movement execution, which may prove instrumental in the development of new non-pharmaceutical strategies in pain management.

## Materials and Methods

To investigate the effects of movement-induced analgesia, we performed five experiments on 130 healthy subjects. Experimental procedures are detailed below, separately for each experiment. All subjects gave their written informed consent and were paid for their participation. The local ethics committee at the Institute of Psychology, Chinese Academy of Sciences approved the procedures.

### Experiment 1

#### Subjects

Behavioral data were collected from 15 healthy volunteers (7 females) aged from 19 to 26 years (M ± SD = 22.1 ± 1.9 years).

#### Nociceptive Stimuli

Nociceptive-specific radiant-heat stimuli were generated by an infrared neodymium yttrium aluminum perovskite (Nd: YAP) laser with a wavelength of 1.34 μm (Electronic Engineering, Italy). At this wavelength, laser pulses directly activate nociceptive terminals in the most superficial skin layers (51). Laser pulses were directed to a circular area (diameter ≈ 4 cm) on the dorsum of subject’s left hand. A He-Ne laser was pointed to the area to be stimulated. The laser beam was then transmitted via an optic fiber with its diameter set at approximately 7 mm (≈ 38 mm^2^) by focusing lenses. The duration of the laser pulse was 4 ms. After each stimulus, the target of the laser beam was shifted by at least 1 cm in a random direction to avoid nociceptor fatigue or sensitization. Subjects were instructed to rate the intensity of pain elicited by each laser stimulus on a numerical rating scale (NRS) ranging from 0 (no sensation) to 10 (the worst pain imaginable), with 4 represents denoting pinprick pain threshold (52). The energy of laser stimuli to be used in the experiment was individually determined by increasing the stimulus energy in steps of 0.25 J, until a rating of 7 out of 10 was obtained, which represents a moderate level of pain. The energies used in the present experiment were 4.0 ± 0.2 J across all subjects, ranging from 3.75 to 4.5 J.

#### Experimental Procedures

Subjects were seated in a comfortable chair in a silent, temperature-controlled room. They were instructed to focus on the laser stimuli, keep their eyes open, and gaze at a fixation on the screen in front of them during the experiment. As illustrated in Fig. 2*A*, the experiment was composed of three blocks of trials, with each block administered under one of the three experimental conditions: control, S1_low, or S1_high. For the control condition, subjects were asked to rest their left hands on a table throughout the whole block. For the S1_low and S1_high conditions, subjects were asked to shake their left hands with a maximal range for 5 s at approximately 1 and 5 Hz, respectively. Each cycle of hand shaking consisted of a rapid motion of the hand away from the body, followed by a return to the original hand position, with minimal whole arm movement. Then, subjects were instructed to rest their left hands on the table after they saw a visual cue of “STOP” appeared in the center of the screen. To ensure that subjects could successfully shake their hands at the required frequency in the S1_low and S1_high conditions, a practice session was included before the formal experiment. During the practice session, subjects were instructed to shake their left hands along with a tracking signal (beeps) presented at either 1 or 5 Hz for a total duration of 5 s in each trial. The beep was a 1000-Hz pure tone with a duration of 50 ms. This phase of the practice session was then followed by a phase in which the tracking signal was removed. Subjects could stop practicing once they could successfully shake their hands at approximately the same frequency without the tracking signal. To rule out the confound introduced by auditory stimuli, no tracking signal was presented during the formal experiment.

For each trial, a fixation was presented once the subjects rested their left hands on the table, and a laser stimulus was delivered on the dorsum of their left hands 1 s after the termination of hand shaking (i.e., the appearance of the fixation on the screen). A visual cue was presented after the laser stimulation prompted the subjects to rate pain intensity (no sensation - the worst pain imaginable) and unpleasantness (no unpleasantness - most unpleasantness) on the same 0-10 NRS. The next trial started 5 s after the subjects responded. Each block contained twelve trials, including two practice trials, which were not used for the following statistical analysis, and ten experimental trials. The order of the three blocks was counterbalanced across subjects, and there was a 10-min break between consecutive blocks.

### Experiment 2

#### Subjects

Behavioral data were collected from 24 healthy volunteers (14 females) aged 19 to 25 years (M ± SD = 21.9 ± 1.7 years).

#### Experimental Procedures

The experiment was composed of three consecutive blocks of trials (Fig. 3*A* & 3*B*). Within each block, data were collected under one of three experimental conditions: control, S1, or S5. For the control condition, subjects were asked to rest their left hands on a table throughout the whole block. For the S1 and S5 conditions, subjects were asked to shake their left hands with a maximal range for 5 s at approximately 5 Hz before resting their left hands on the table. A practice session with a tracking signal at 5 Hz was included before the formal experiment to ensure that subjects could successfully shake their hands at the required frequency. For each trial, a fixation was presented once the subjects rested their left hands on the table, and a laser stimulus was delivered on the dorsum of their left hands 1 or 5 s after the termination of hand shaking, respectively. The energies of laser stimuli were 4.0 ± 0.3 J across all subjects, ranging from 3.5 to 4.5 J in the present experiment. The rest of experimental procedures were identical to that of Experiment 1.

### Experiment 3

#### Subjects

Behavioral data were collected from 25 healthy volunteers (14 females) aged 19 to 27 years (M ± SD = 22.5 ± 2.3 years).

#### Experimental Procedures

The experiment was composed of five blocks of trials, each administered under one of five experimental conditions: control, S1, S15, S20, or S30 (Fig. 3*A* & 3*C*). For the S1, S15, S20, and S30 conditions, subjects were asked to shake their left hands with a maximal range for 5 s at approximately 5 Hz before resting their left hands on the table. A laser stimulus was delivered on the dorsum of their left hands 1, 15, 20, or 30 s after the termination of hand shaking, respectively. The energies of laser stimuli were 4.2 ± 0.3 J across all subjects, ranging from 3.75 to 4.5 J in the present experiment. The rest of experimental procedures were identical to that of Experiment 1.

### Experiment 4

#### Subjects

Behavioral data were collected from 25 healthy volunteers (14 females) aged 19 to 28 years (M ± SD = 23.6 ± 2.4 years).

#### Experimental Procedures

The experiment was composed of five blocks of trials, each administered under one of five experimental conditions: control, S1, S10, CS1, or CS10 (Fig. 3*A* & 3*D*). For the S1 and S10 conditions, subjects were asked to shake their left hands with a maximal range for 5 s at approximately 5 Hz before resting their left hands on the table. A laser stimulus was delivered on the dorsum of their left hands 1 or 10 s after the termination of hand shaking, respectively. For the CS1 and CS10 conditions, subjects were asked to shake their right hands with a maximal range for 5 s at approximately 5 Hz, and a laser stimulus was delivered on the dorsum of their left hands 1 or 10 s after the termination of hand shaking, respectively. The energies of laser stimuli were 4.2 ± 0.4 J across all subjects, ranging from 3.0 to 4.5 J in the present experiment. The rest of experimental procedures were identical to that of Experiment 1.

### Experiment 5

#### Subjects

Behavioral and EEG data were collected from 41 healthy volunteers (24 females) aged 20 to 27 years (M ± SD = 22.7 ± 1.9 years).

#### Experimental Procedures

The experiment was composed of five blocks of trials, each administered under one of five experimental conditions: control, S1, S15, CS1, or CS15 (Fig. 3*A* & 3*E*). The experimental procedures were identical to those of Experiment 4, except that instead of the S10 and CS10 conditions, the S15 and CS15 conditions were included, in which the temporal delay between the termination of hand shaking and laser stimulus was 15 s in both conditions. In addition, a visual cue was presented 5 s after each laser stimulus prompted the subjects to rate pain intensity and unpleasantness on the same 0-10 NRS. The energies of laser stimuli were 3.5 ± 0.4 J across all subjects, ranging from 3 to 4.5 J in the present experiment.

#### EEG Recording

EEG data were recorded using 64 Ag-AgCl scalp electrodes placed according to the International 10-20 system (Brain Products GmbH; pass band: 0.01-100 Hz; sampling rate: 1000 Hz). The nose was used as reference, and electrode impedances were kept lower than 10 kΩ. To monitor ocular movements and eye blinks, electrooculographic signals were simultaneously recorded from two electrodes: one placed over the lower eyelid of the left eye, the other placed 1 cm lateral to the outer corner of the left orbit.

#### EEG Preprocessing

EEG data were processed using EEGLAB (53), an open-source toolbox running in the MATLAB (Mathworks, USA) environment. Continuous EEG data were band-pass filtered between 1 and 30 Hz. EEG epochs were extracted using a window analysis time of 3000 ms (1000 ms before to 2000 ms after the onset of laser stimuli) and baseline-corrected using the prestimulus interval. Trials contaminated by eye-blinks and movements were corrected using an independent component analysis algorithm (53).

#### Time-domain Analysis

EEG epochs in each condition were averaged in the time domain, thus yielding five average waveforms time-locked to laser stimuli for each subject. The N1, N2, and P2 amplitudes were measured from the averaged LEP waveform of each subject. The N2 and P2 waves were defined as the most negative and positive deflections between 150 and 500 ms after laser stimulation at the central electrode (Cz-nose), respectively (26). The N1 wave was defined as the most negative deflection preceding the N2 wave, which showed a contralateral central scalp distribution. A bipolar montage, i.e., central electrode contralateral to the stimulated side referenced to Fz (C4-Fz), was adopted to optimally detect the N1 wave (54, 55). Single-subject average waveforms were subsequently averaged across subjects to obtain group-level waveforms. Group-level scalp topographies of these LEP waves were computed by spline interpolation.

#### Prestimulus EEG oscillations

To assess the modulation of voluntary movement of spontaneous EEG oscillations, we extracted prestimulus EEG signals from a time window ranging from −1000 to 0 ms relative to the laser stimulus onset. For each subject, prestimulus EEG signals were transformed to the frequency domain using a discrete Fourier transform, yielding an EEG spectrum ranging from 1 to 30 Hz. Single-subject EEG spectra were averaged across subjects to obtain group-level prestimulus EEG spectra for each experimental condition. Since prestimulus alpha oscillations have been shown to be modulated by hand movement and somatosensory feedback (43), we tested the *a priori* hypothesis that the modulation of voluntary movement on pain perception was influenced by prestimulus alpha oscillations. Specifically, spectral amplitude of alpha oscillations was extracted at 8-12 Hz from bilateral centro-parietal regions (i.e., C3, CP3, C4, and CP4) for the following statistical analysis.

To localized the sources of spontaneous alpha oscillations, we used a beamforming algorithm known as dynamic imaging of coherent sources (56), implemented in FieldTrip (57). This algorithm computes a spatial filter based on a leadfield matrix and a cross-spectral density matrix. The leadfield matrix, containing coefficients that reflects the mapping between current sources and scalp potential differences, was computed for a three-dimensional grid with a 1-cm resolution, using a realistically shaped three-shell boundary-element volume conduction model based on the Montreal Neurological Institute template brain. The cross-spectral density matrix was computed for alpha oscillations (i.e., 8-12 Hz) using a multitaper frequency transformation. The estimated power in source space yielded an estimate of the neural sources of spontaneous alpha oscillations for each experimental condition.

### Statistical analysis

For all experiments, subjective ratings of pain intensity and unpleasantness were compared using a one-way repeated-measures analysis of variance (ANOVA) with a within-subject factor of condition. Greenhouse-Geisser adjustments were used in light of violations of sphericity. When the main effect was significant (*P* < 0.05), post hoc paired-sample t-tests were performed, and Bonferroni-correction was applied for multiple comparisons. Partial eta-squared (*η*_*p*_^*2*^) was calculated to reflect the effect size. For Experiment 5, the same statistical analyses were conducted on the N1, N2, and P2 amplitudes, as well as the spectral amplitude of prestimulus alpha oscillations. To demonstrate the brain regions with movement-induced changes of spontaneous alpha oscillations, a point-by-point one-way repeated-measures ANOVA was performed for each electrode at the scalp level and for each grid point at the source level. This procedure yielded a scalp topography and a source distribution of *F* values representing the significance level of the modulation of voluntary movement on alpha oscillations, respectively. To account for the multiple comparisons, a false discovery rate procedure was performed.

## Acknowledgments

This research was supported by the National Natural Science Foundation of China (No. 31822025, 31701000, 31671141).

## SI Appendix

**Table S1.** Subjective ratings (Mean ± SD) of pain intensity and unpleasantness for each experimental condition (S1_low, S1_high, and Control) in Experiment 1.

|  | S1_low | S1_high | Control |
| --- | --- | --- | --- |
| Pain intensity | 6.70 ± 1.17 | 6.11 ± 1.50 | 7.09 ± 1.36 |
| Unpleasantness | 6.20 ± 1.71 | 5.42 ± 1.82 | 6.52 ± 1.65 |

**Table S2.**
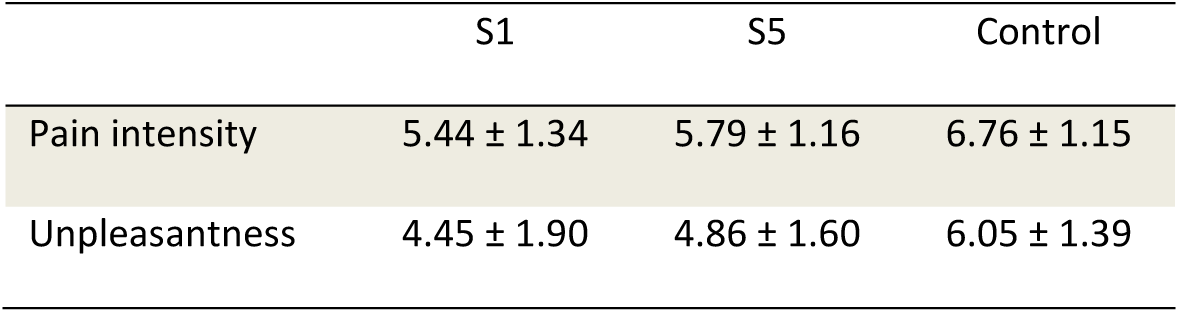
Subjective ratings (M ± SD) of pain intensity and unpleasantness for each experimental condition (S1, S5, and Control) in Experiment 2.

|  | S1 | S5 | Control |
| --- | --- | --- | --- |
| Pain intensity | 5.44 ± 1.34 | 5.79 ± 1.16 | 6.76 ± 1.15 |
| Unpleasantness | 4.45 ± 1.90 | 4.86 ± 1.60 | 6.05 ± 1.39 |

**Table S3.**
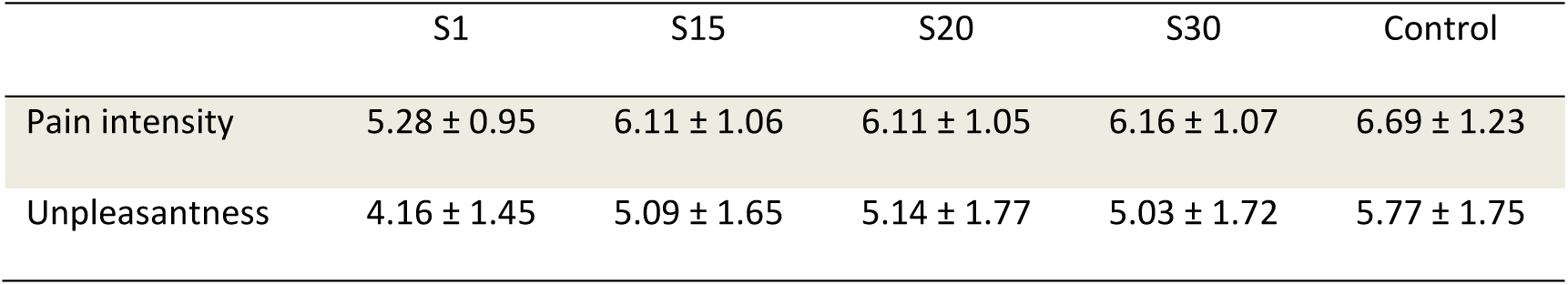
Subjective ratings (M ± SD) of pain intensity and unpleasantness for each experimental condition (S1, S15, S20, S30, and Control) in Experiment 3.

**Table S4.** Subjective ratings (M ± SD) of pain intensity and unpleasantness for each experimental condition (S1, S10, CS1, CS10, and Control) in Experiment 4.

|  | S1 | S10 | CS1 | CS10 | Control |
| --- | --- | --- | --- | --- | --- |
| Pain intensity | 5.01 ± 1.23 | 5.61 ± 1.26 | 6.20 ± 1.06 | 6.57 ± 1.00 | 6.59 ± 1.08 |
| Unpleasantness | 3.62 ± 1.96 | 4.03 ± 2.21 | 4.60 ± 1.91 | 5.11 ± 1.93 | 5.32 ± 1.97 |

**Table S5.** Subjective ratings (M ± SD) of pain intensity and unpleasantness, N1, N2, and P2 amplitudes, as well as the spectral amplitude of prestimulus alpha oscillations (Pre-α) for each experimental condition (S1, S15, CS1, CS15, and Control) in Experiment 5.

|  | S1 | S15 | CS1 | CS15 | Control |
| --- | --- | --- | --- | --- | --- |
| Pain intensity | $4.51 \pm 1.03$ | $5.36 \pm 0.93$ | $5.98 \pm 0.84$ | $5.97 \pm 1.07$ | $6.31 \pm 1.02$ |
| Unpleasantness | $3.38 \pm 1.78$ | $4.27 \pm 1.86$ | $4.86 \pm 1.79$ | $4.92 \pm 1.91$ | $5.38 \pm 1.85$ |
| N1 amplitude ($\mu V$) | $-2.93 \pm 3.61$ | $-4.81 \pm 4.87$ | $-4.75 \pm 4.53$ | $-5.45 \pm 5.51$ | $-5.24 \pm 5.09$ |
| N2 amplitude ($\mu V$) | $-7.15 \pm 7.60$ | $-16.20 \pm 12.04$ | $-12.38 \pm 12.76$ | $-16.71 \pm 14.39$ | $-17.40 \pm 15.36$ |
| P2 amplitude ($\mu V$) | $9.30 \pm 10.88$ | $15.41 \pm 13.41$ | $15.57 \pm 11.71$ | $19.00 \pm 11.57$ | $19.80 \pm 14.30$ |
| Pre-α amplitude ($\mu V$) | $-0.03 \pm 0.27$ | $0.26 \pm 0.47$ | $0.01 \pm 0.21$ | $0.23 \pm 0.39$ | $0.23 \pm 0.36$ |

